# Reassessment of Mendelian gene pathogenicity using 7,855 cardiomyopathy cases and 60,706 reference samples

**DOI:** 10.1101/041111

**Authors:** Roddy Walsh, Kate L Thomson, James S Ware, Birgit H Funke, Jessica Woodley, Karen J McGuire, Francesco Mazzarotto, Edward Blair, Anneke Seller, Jenny C Taylor, Eric V Minikel, Exome Aggregation Consortium, Daniel G MacArthur, Martin Farrall, Stuart A Cook, Hugh C Watkins

## Abstract

The accurate interpretation of variation in Mendelian disease genes has lagged behind data generation as sequencing has become increasingly accessible. Ongoing large sequencing efforts present huge interpretive challenges, but also provide an invaluable opportunity to characterize the spectrum and importance of rare variation. Here we analyze sequence data from 7,855 clinical cardiomyopathy cases and 60,706 ExAC reference samples to better understand genetic variation in a representative autosomal dominant disorder. We show that in some genes previously reported as important causes of a given cardiomyopathy, rare variation is not clinically informative and there is a high likelihood of false positive interpretation. By contrast, in other genes, we find that diagnostic laboratories may be overly conservative when assessing variant pathogenicity. We outline improved interpretation approaches for specific genes and variant classes and propose that these will increase the clinical utility of testing across a range of Mendelian diseases.

**One Sentence Summary:** Comparing the frequency of very rare variation between patient cohorts and very large genomic reference datasets enables the reliable re-evaluation of genes previously implicated in Mendelian disease and more accurate assessment of the likely pathogenicity of different classes of variants.

## Introduction

Interpretation of rare genetic variation, whether in a clinical diagnostic or research setting, has not kept pace with accelerating data generation using high-throughput DNA sequencing. Increasingly extensive gene panels are used to interrogate the growing number of genes implicated in Mendelian diseases *(1)*. However, such panels only modestly increase the number of high-confidence diagnostic results while identifying ever larger numbers of variants of uncertain significance (VUS) *(2–4)*. The likelihood that a rare variant is indeed pathogenic depends on the pre-test probability that the patient has the relevant disease. Thus testing of a wide panel of genes for loosely related phenotypes is expected to yield an even higher proportion of false positive variants. Analysis of all genes in the genome regardless of clinical indication, and of putative pathogenic variants found as ‘incidental findings’, will exacerbate the problem further. These concerns have required adoption of conservative interpretation guidelines *(5)* potentially limiting the true utility of genetic testing.

Central to the challenge of rare variant interpretation is the paradox that individually rare variants are now seen to be collectively common. While it is accepted that a common variant can be excluded as a cause of a rare and penetrant Mendelian disease, the community has been slower to recognise that many rare variants identified in Mendelian disease genes are innocent bystanders, and some ‘rare’ variants are not rare at all. Recent population sequencing efforts have raised awareness of these issues (1000 Genomes Project *(6, 7)*, Exome Sequencing Project http://evs.gs.washington.edu/EVS) but the full extent is now revealed in the Exome Aggregation Consortium (ExAC) study (http://exac.broadinstitute.org) where the average ExAC exome contains 7.6 rare non-synonymous variants (minor allele frequency [MAF] <0.1%) in well-characterized dominant disease genes, the large majority being very rare, or ‘private *(8)*. Clearly only a small minority can actually cause a penetrant Mendelian disease *(9)*.

The challenges of variant interpretation in Mendelian disorders are particularly well illustrated by inherited cardiomyopathies: Hypertrophic Cardiomyopathy (HCM), Dilated Cardiomyopathy (DCM) and Arrhythmogenic Right Ventricular Cardiomyopathy (ARVC). These largely autosomal dominant disorders are relatively common, genetically heterogeneous and medically important *(10)* and consequently cardiomyopathy genes feature prominently in the American College of Genetics and Genomics (ACMG) list of proposed genes to be routinely analysed in all exome or genome sequencing *(11)*. While clinical genetic testing in cardiomyopathy has been available for over a decade, the number of genes reported as disease-causing has increased dramatically in recent years, often without robust evidence.

Here we leverage two very substantial resources to better understand and interpret rare variation in cardiomyopathy genes, thereby illustrating the scale of the challenges but also possible solutions. We aggregated sequence data for cardiomyopathy genes from 7855 individuals with a diagnosis of cardiomyopathy, sequenced in accredited diagnostic laboratories with clinical-grade variant interpretation (4,187 individuals sequenced at Oxford Medical Genetics Laboratories [OMGL] and reported here for the first time, and a recently published series of 3,668 individuals from the Partners HealthCare Laboratory of Molecular Medicine [LMM] *(4, 12)*). These data were compared with sequence data from 60,706 reference samples from the ExAC consortium. ExAC cohorts likely to be enriched for Mendelian disease were not included in this dataset and while cardiomyopathy has not been excluded in ExAC cases, the selection criteria for the component cohorts would not be expected to enrich for inherited cardiac conditions. Of crucial importance, the ExAC resource is the first dataset powered to assess variant alleles present in the population at a range of 1:1,000-100,000 that might previously have been considered pathogenic yet may in fact be too common to cause penetrant Mendelian disease.

By empirically deriving an upper bound for the allele frequency of confirmed cardiomyopathy-causing mutations and comparing the number of variants below this threshold in cases and in ExAC, we show that for some of the genes included in cardiomyopathy panel tests there is little or no excess of rare variants in cases and hence such variation cannot be interpreted. Research studies that have included such variants, or tested genes unrelated to the presenting condition, have reached erroneous conclusions. In contrast, our data suggest that a large fraction of variants found in validated genes that are currently reported as VUSs by diagnostic laboratories are actually likely to be pathogenic. Comparison of the localisation of rare variants in cases with findings from very large numbers of controls also yielded valuable protein domain-specific information. Hence, through these types of analyses it is possible to define the genes, regions of genes and/or classes of variants that can be reliably interpreted in a clinical setting and in doing so, increase clinical diagnostic yields. These approaches, when combined with disease priors, promise to enhance our interpretation of rare genetic variation, thus maximising the utility of sequencing for precision medicine.

## Results

### Deriving allele frequencies for potentially deleterious variants

Of the three cardiomyopathies, HCM is the most prevalent (affecting 1 in 500*(13)*) and HCM genes are known to harbour recurrent disease-causing variants. The single most common confirmed pathogenic variant, in both clinical cohorts, was *MYBPC3* c.1504C>T (p.Arg502Trp), found in 104/6179 HCM cases (1.7%, 95CI 1.4-2.0%); this variant was only observed 3 times in ExAC (MAF 2.5×10^−5^). We therefore applied a minor allele frequency (MAF) cut-off of 1×10^−4^ as a conservative upper-bound as variants more frequent than this in the general population would not be expected to be pathogenic (see Supplementary Note 1). This MAF does not exclude more common deleterious founder variants in specific populations where the genetic architecture of cardiomyopathy is not well defined.

### Comparison of disease cohort variants with ExAC variants (ExAC MAF < 1×10^−4^)

Variants identified by sequencing of putative cardiomyopathy genes in cases (n=7855) were collated by disease and gene (Supplementary Tables S1A and S1B). There were no significant differences in rare variant frequency between the two clinical genetics laboratories, so data were combined (Supplementary Tables S3A and S3B). We compared the burden of rare protein altering variants (ExAC MAF < 1×10^−4^) detected in 20 HCM genes, 48 DCM genes and 8 ARVC genes in HCM, DCM and ARVC cases, respectively, with the burden observed in ExAC. Predicted truncating (nonsense, frameshift, or canonical splice site) and non-truncating (missense or small in-frame deletions) variants were analysed separately. (Supplementary Tables S4A, S4B and S4C).

As expected *(14–16)*, rare variation in the two major HCM genes accounted for the majority of observed variation in HCM cases (*MYBPC3*, 19.0% of cases; *MYH7*, 14.2%). Rare variants were less numerous in other well-characterised HCM genes (*TNNI3, TNNT2, TPM1, MYL2, MYL3, ACTC1, PLN*) and phenocopy genes (*GLA, LAMP2, PRKAG2*) (≤2% cases per gene). For each of these genes there is a significant (P<0.05 after Bonferroni correction) excess of variation in cases as compared with ExAC, confirming their association with disease (Fig. 1 and Supplementary Table S5A). However, for some more recently reported HCM genes (*TNNC1(17)*, *MYOZ2(18)*, *ACTN2(19), ANKRD1(20)*) there was no significant excess of rare genetic variation in these HCM cases.

**Figure 1:**
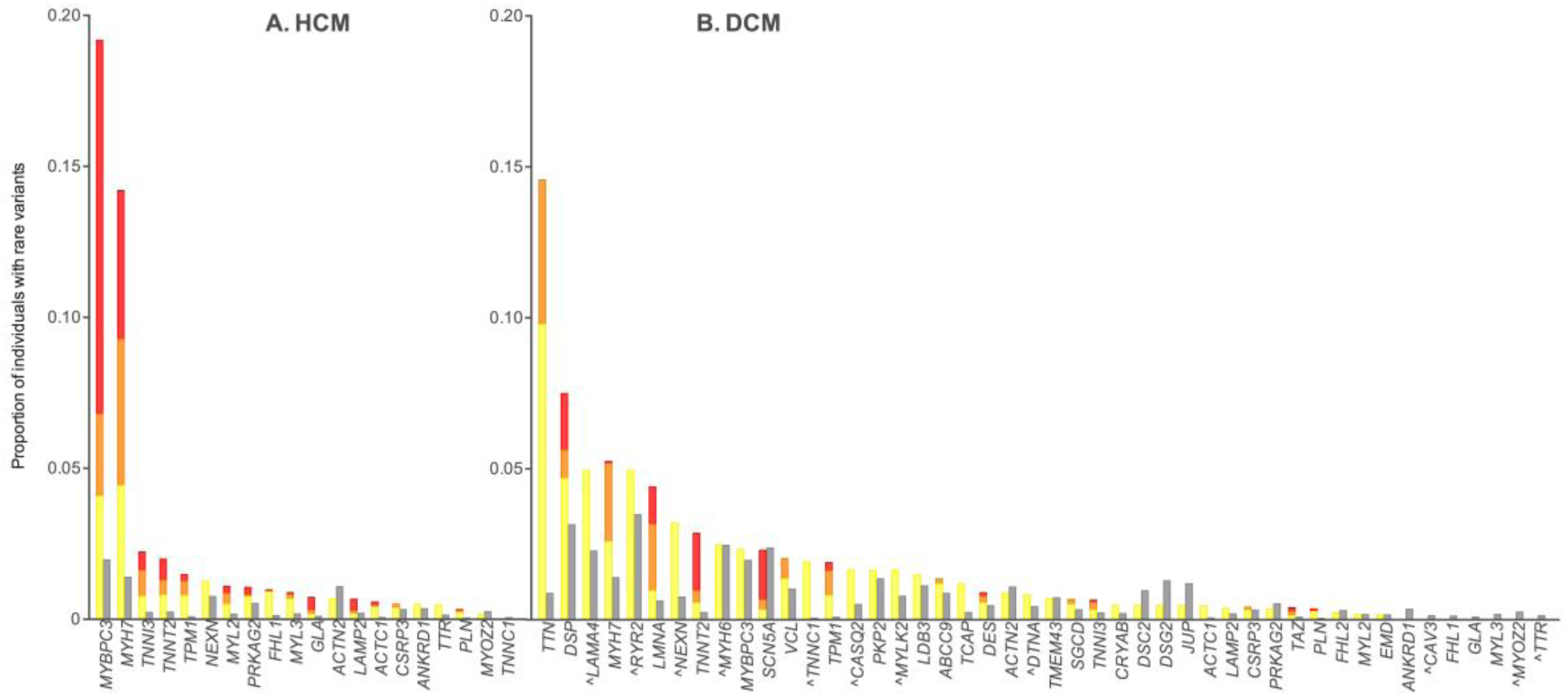
Proportion of individuals with rare variants in HCM and DCM in combined clinical cohorts (data from OMGL and LMM) compared to ExAC (grey columns). Variants counted were single nucleotide change or small insertion/deletion variant detected in the coding region ±2bp, with an ExAC minor allele frequency (MAF) of <1×10^−4^ (see Methods). **Clinical cohorts**: HCM n=632 to 6179, DCM n=121 to 1315 (see Supplemental tables S5A and S5B). Overlaid is information on reported pathogenicity class (red = pathogenic (P), orange = likely pathogenic (LP), yellow = variant of uncertain significance (VUS). See Supplemental Tables S4A and S4B for full details. ^ = genes analyzed in fewer than 200 cases. **ExAC:** n=mean of total adjusted allele count for rare variant carriers. For HCM genes n ranged from 47153 to 60647, for DCM genes n was 42697 to 60647 (see Supplemental tables S5A and S5B). CTF1 and RBM20 were removed from analysis due to poor coverage in ExAC.

DCM is reportedly highly genetically heterogeneous with up to ~60 genes previously implicated *(16, 21, 22)*. In the clinical cohorts, truncating variants in *TTN* were most common (14.6%) in keeping with our findings in large research cohorts *(23, 24)*. The prevalence of rare variants in other well-characterised DCM genes was modest (*MYH7* 5.3%, *LMNA* 4.4%, *TNNT2* 2.9% and *TPM1* 1.9%) but significantly enriched as compared to ExAC (Fig. 1 and Supplementary Table S5B). However, with the exception of truncating variants in *DSP* (2.8%), there was limited burden and modest or no significant excess variation in the remaining 40 genes tested. In ARVC, the five major genes each showed significant excess in cases (Supplementary Table S5C).

Overall, the yield of pathogenic (P) and likely pathogenic (LP) variants was 32% in HCM, 13% in DCM (but note that *TTN* was only sequenced in a third of samples) and 36% in ARVC. Of note, however, even variants of uncertain significance (VUS) were seen in substantial excess over controls for all of the genes robustly supported by pathogenic and likely pathogenic variants, suggesting that clinical labs may be overly conservative.

To assess for any confounding effects of population stratification between cases and controls, the LMM DCM cohort (the only cardiomyopathy dataset for which individual, self-reported ethnicity data was available) was compared to ExAC across all case and control samples and separately for “white” cases versus the non-Finnish European subset of ExAC. There was full correlation between the genes that were significantly enriched in both comparisons (Supplementary Table S13), with a Pearson correlation coefficient of 0.97 between the case excess values for genes in the two comparisons (Supplementary Fig 1), showing that population stratification effects are not a confounder in this study.

### Using case: control variant burdens to interpret individual variants

Many variants in confirmed disease-genes can be interpreted with confidence based on cumulative experience (e.g. segregation in families, multiple occurrences in cases, *de novo* mutations) and/or functional insights (e.g. null alleles in haploinsufficient genes), and this underlies current clinical utility. But our ability to evaluate the pathogenicity of the large number of variants that are seen for the first time depends on the signal to noise ratio. For each gene and variant class we calculated two related metrics: the odds ratio (OR) (ratio of odds of cardiomyopathy comparing rare variant carriers with non-carriers) and the etiological fraction (EF), a commonly used measure in epidemiology *(25–27)* which estimates the proportion of cases in which the exposure (in this case a rare variant in a gene) was causal (see Supplementary Note 2).

These analyses reaffirm high ORs and EFs in key cardiomyopathy genes but also highlight a number of previously reported cardiomyopathy genes that show limited disease association when compared with a very large number of controls (Fig. 2 and Supplementary Tables S5A, S5B and S5C). As expected, many genes have divergent results for truncating as compared to non-truncating variants. *MYH7* is a typical example, with an OR of 1 [0.5-4.5] for truncating variants vs. an OR of 12 [10.9-13.3] for non-truncating variants in HCM cases.

**Figure 2:**
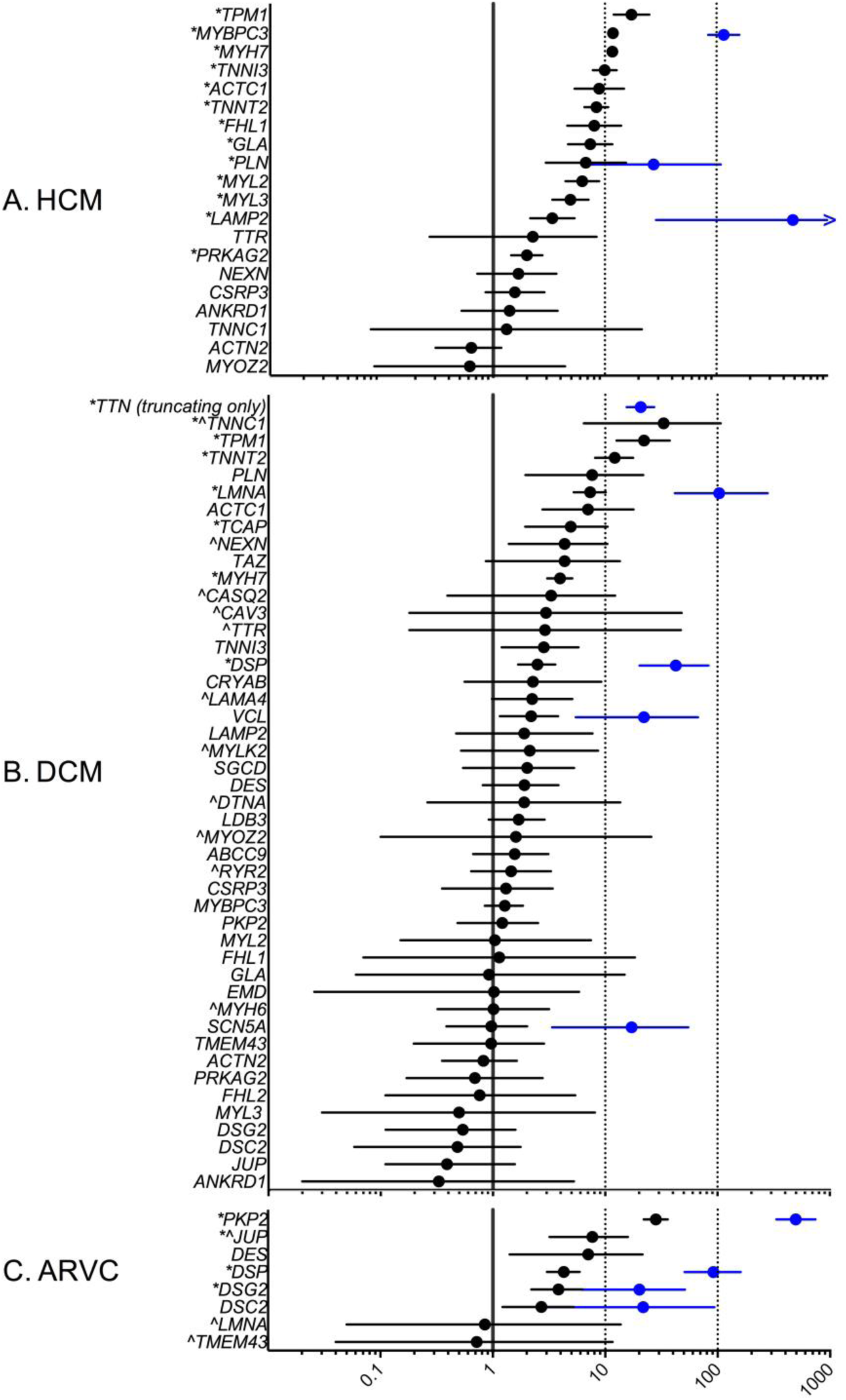
Odds ratios (OR) with 95% CI for each gene tested in the HCM (n=632 to 6179), DCM (n=121 to 131) and ARVC (n=93 to 361) clinical cohorts compared to ExAC reference samples (n=mean of total adjusted allele count for rare variant carriers. For HCM genes n=47153 to 60647, DCM genes n=42697 to 60647 and ARVC genes n=51126 to 60218). Data has been plotted (log_10_ scale) for all protein-altering variants (black) and truncating variants (blue). For truncating variants, OR with 95% CI have been plotted on for genes where a statistically significant difference was observed for this variant type on FET. *= Statistically significant Fishers exact test (FET) (P=0.05 with Bonferroni correction, HCM P=<0.0025, DCM P=0.001 and ARVC P=<0.006.) See Supplemental tables S5A, S5B and S5C for data use to generate this plot. ^ =genes analyzed in fewer than 200 cases. CTF1 and RBM20 were removed from analysis due to poor coverage in ExAC.

This observation confirms the widely accepted view that missense alleles of *MYH7* act as dominant negatives in HCM whereas truncating variants are not pathogenic. In genes where truncating alleles are disease-causing, ORs are typically higher due to the lower rate of truncating variants in the population. As expected, truncating variation in *MYBPC3* associates strongly with HCM (the result of haploinsufficiency *(28)*), but neither truncating nor non-truncating variation in *MYBPC3* shows significant association with DCM (OR=1.3 [0.8-1.8]; EF=0.21 [0-0.46]), a finding that fits with mechanistic insights but challenges some widely held viewpoints *(29, 30)*. Amongst the ARVC genes, truncating alleles are informative for four major genes (and particularly common for *PKP2* and *DSP*), whereas non-truncating variants in these genes are difficult to interpret reliably (Fig. 2 and Supplementary Tables S4C and S5C).

### Using protein domain knowledge to improve variant interpretation

At the gene-level, ORs for non-truncating variants are typically modest, and in the absence of prior clinical experience or functional data, interpretation is often uncertain. This may be improved by considering protein topology, as pathogenic variants often cluster in specific regions in cases *(31, 32)*. We evaluated the distribution of rare missense alleles in *MYH7*, which encodes a protein with well-characterised functional and structural domains, to assess whether adding a systematic analysis of variant distribution refines such interpretation. Non-random mutation cluster analysis *(33)* revealed a significant cluster (p<3×10^−15^, False Discovery Rate (FDR) q<5×10^−13^) between residues 181 and 937 in HCM cases, whereas variants in ExAC controls were depleted in this region and instead clustered between residues 1271 and 1903 (p<3×10^−8^, FDR q<4×10^−5^) (Fig. 3). These data more precisely define the boundaries of mutation-enriched and depleted zones that can be used to generate more discriminating EFs, e.g. for rare variants in HCM patients, EFs range from 0.97 in the HCM cluster to 0.67 in the control cluster (Figs. 3 and 4).

**Figure 3:**
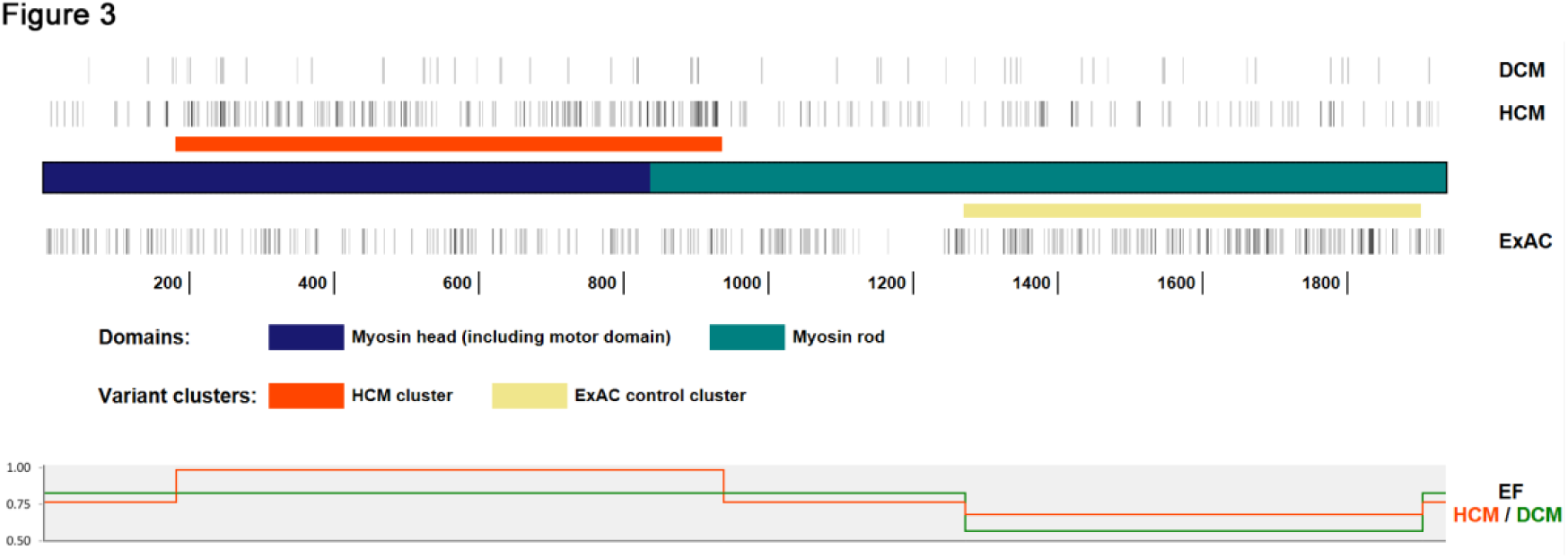
Distribution of rare (ExAC MAF <1×10-4) MYH7 missense variants in HCM (n=864) and DCM (n=69) clinical cohorts and ExAC controls (n=816), with the key myosin protein regions highlighted. Using non-random mutation cluster analysis (NMC), (refer to Online Methods), a significant variant cluster (p<3×10-15, False Discovery Rate (FDR) q<5×10-13) was identified between residues 181 and 937 (involving the motor domain, leverarm and part of the rod) in HCM cases, and depletion in this region and a significant cluster (p<3×10--8, FDR q<4×10-5) between residues 1271 and 1903 (in the part of the rod that forms the filament backbone) in controls. The etiological fraction (EF) for a rare MYH7 missense variant identified in a HCM proband ranges from 0.97 in the HCM cluster to 0.67 in the control cluster. Vertical grey bars depict the positions of variants in cohorts, grey scale showing variant density where variants are co-incident.

**Figure 4:**
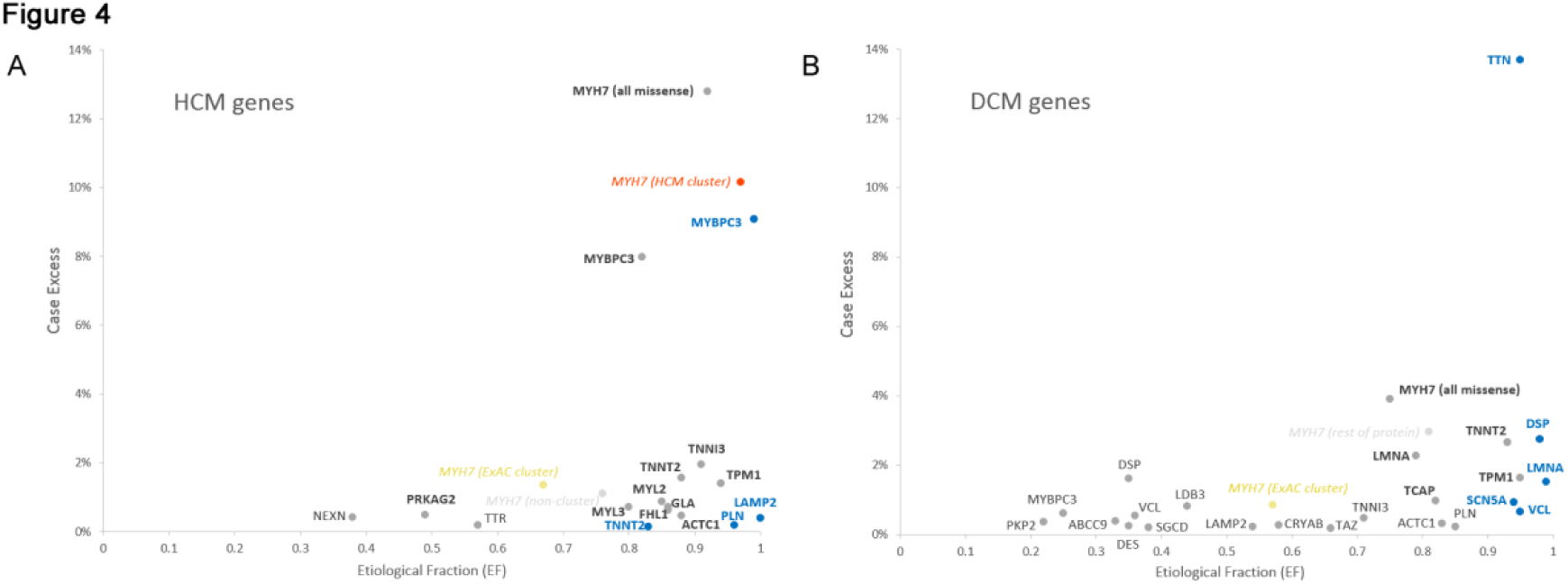
An overview of the genetic landscape of HCM (**A**) and DCM (**B**) for truncating (blue) and non-truncating (grey) variants, as well as *MYH7* missense variants in the clusters identified in Fig. 3 (orange, disease cluster; yellow, ExAC control cluster). The case excess (y-axis) is the frequency of rare variation in disease cohorts over and above the frequency in ExAC controls and indicates the relative importance of the gene and variant class to the genetic etiology of each cardiomyopathies. The etiological fraction (EF) (x-axis) is a measure of the interpretability of variants of this class, being an estimate of the proportion of affected carriers where the variant caused the disease (see Supplementary Tables S4A and 4B for full details). This measure is an average of all variants of a given class, some of which will be pathogenic but others benign.

### Application of findings

To facilitate the application of these findings for research and clinical use we provide an overview of the genetic landscape of cardiomyopathy as represented by patient referrals received by UK and US clinical testing laboratories. This shows both the relative importance of cardiomyopathy genes within these patient populations (measured as a ‘case excess’), and their interpretability (expressed as the etiological fraction) (Fig. 4). Furthermore, we have created a web resource, Atlas of Cardiac Genetic Variation (http://cardiodb.org/ACGV) to provide easy access to our data to aid those assessing the relevance of specific genes and classes of variant to cardiomyopathies. We believe this provides an exemplar of how large case series combined with ExAC control data can be used across diseases to refine our understanding of disease genetics.

### Reassessing extended gene panel studies of cardiomyopathy

A number of recent research studies *(29, 30, 34, 35)* using extended gene panels have reported putative genetic overlap between diverse inherited cardiac diseases that poses great challenges for clinical interpretation and appears at odds with known disease mechanisms. We surmise that many such studies have not adequately accounted for background genetic variation, have relied on variant data from incompletely annotated disease-centred databases and have not used segregation. Here, using ExAC control data, we present a reanalysis of two of these research studies *(30, 34–36)*. In the research HCM cases, the excess variation in the known HCM genes (e.g. *MYBPC3*, *MYH7*) is substantial. In stark contrast, the measured variation in DCM, ARVC and ion channel genes in the HCM patients, while also substantial, is similar to that seen in ExAC samples with little, if any, excess burden (Fig. 5A and Supplementary Table S6). This suggests that the large majority of these variants, though individually rare, are benign bystanders, and that any overlap between the disorders has been over-estimated. In the DCM research studies, even though sequencing was limited to putative DCM genes, some genes which were proposed on the basis of these and other recent studies as amongst the most common causes DCM, such as *MYBPC3*, *MYH6* and *SCN5A (29, 30, 35)* in fact showed no excess variation (Fig. 5B and Supplementary Table S7).

**Figure 5:**
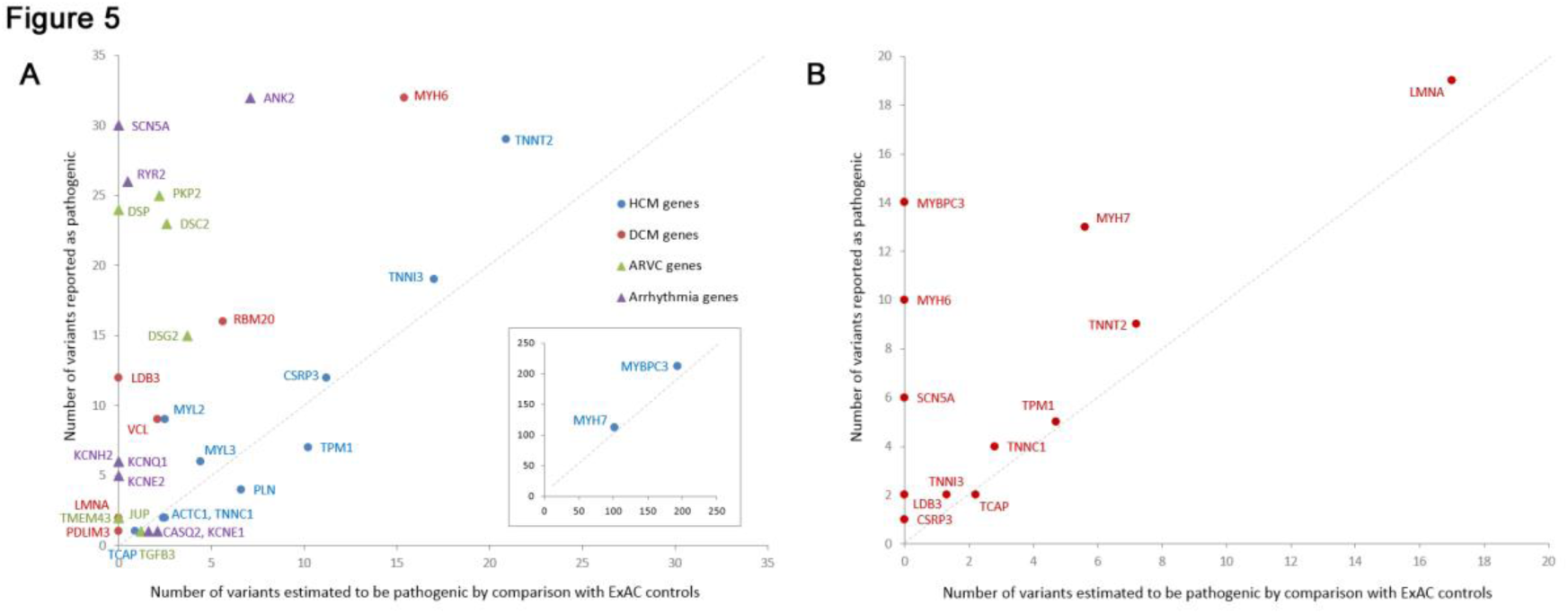
Comparison of the number of variants reported as putatively pathogenic in HCM (**A**) and DCM (**B**) in research studies (using generic analysis criteria such as variant class, missense effect predictions and variant population frequency in ESP) with those predicted as pathogenic by the excess of variation in cases over ExAC controls in each gene. For the HCM study *(34)* (**A**), genes are colored by the cardiac disease for which they are primarily associated, as defined by Lopes et al *(34)*. While there is good concordance between the research findings and the ExAC predictions for established HCM genes, for genes primarily associated with DCM, ARVC and arrhythmias, the variation in cases is similar to that in controls. In the DCM study *(30, 35, 36)* (**B**), variation burden in MYBPC3, SCN5A and MYH6 is similar between the published research cases and ExAC controls, suggesting most variants in these genes are unlikely to be causing DCM.

### Reclassification of variants previously reported as pathogenic using ExAC

We examined the allele frequency of variants previously reported to cause HCM, DCM or ARVC, as catalogued by HGMD (Supplementary Tables S8-S10). A substantial number of purported disease-causing variants for HCM (25.2%, 322/1280), DCM (29.2%, 222/759) and ARVC (34.6%, 167/483) were observed in ExAC. While presence in ExAC does not preclude pathogenicity, a significant number are present at an allele frequency incompatible with causation of penetrant cardiomyopathy (6.5% of HCM, 11.9% of DCM and 13.5% of ARVC variants are present at MAF >1×10^−4^, Supplementary Table S11). Tellingly, some 75% of HGMD variants that could not be excluded as disease-causing using the NHLBI Exome Sequencing Project (the largest control dataset prior to ExAC), due to a control allele count of one, can be discounted by ExAC refinement. In total, 11.7%, 19.6% and 20.1% of individuals in ExAC have reported HCM, DCM and ARVC variants respectively, far in excess of disease prevalence. Hence variant prioritization based on HGMD status alone is not advised for cardiomyopathy genes, a fact that is increasingly apparent with larger control datasets.

## Discussion

We present an analysis of data from 7,855 individuals referred for clinical sequencing and variant interpretation for inherited cardiomyopathies, alongside 60,706 ExAC reference samples. These data exemplify the many challenges of variant interpretation in genetically heterogeneous disorders. We propose that in the absence of large matched case control series, the approaches described here, using data from large patient cohorts and broader reference datasets such as ExAC, may be applied to a range of multi-genic, multi-allelic diseases

We show that the pathogenicity of disease genes originally identified through family linkage are resoundingly validated, for example the majority of sarcomere genes in HCM. However, genes implicated in cardiomyopathy through candidate gene studies, including genes on panel tests in routine clinical use, are often not convincingly associated with disease. For example, *MYBPC3*, *MYH6* and *SCN5A* have all been reported to be major contributors to DCM *(29, 30, 35)*, but show little or no excess burden despite adequate numbers and power; instead we see that these are in fact genes that have the highest background variation.

We also show that it is crucial not only to distinguish variant classes, but to assess these in light of known disease mechanisms for each gene and disorder. For example, cardiomyopathy-causing variants in most myofilament proteins incorporate into the sarcomere and act as dominant negatives (HCM mutations are activating, whereas DCM mutations depress myofibrillar function) *(37)*. Hence protein-truncating variants, which do not incorporate would not be expected to cause these conditions, and this is borne out in our data. In contrast, *MYBPC3* truncation alleles cause HCM through haploinsufficiency, making it unlikely that they could also cause DCM, which we confirm with our findings.

We summarize our analyses of cardiomyopathy genes in two measures, capturing the contribution of each gene to a disease (case excess) and our ability to interpret variation in each gene (etiological fraction). Etiological fraction (EF) can be interpreted as the proportion of affected carriers where the variant caused the disease, i.e. the proportion of true positives. EF is based on pooled rare variant frequency data so it summarizes the average risk across many variants in a gene (some of which will be pathogenic but others benign); EF will be particularly useful for selecting panels of genes that are informative for discrete phenotypes. Of critical importance, the probability that a novel variant is pathogenic depends on the clinical status of the individual carrying the variant. Thus when variants are found in individuals with a remote/unrelated clinical diagnosis, or as an incidental finding during exome or genome sequencing, the proportion of variants expected to be pathogenic will be considerably lower.

While detailed phenotyping of the cardiomyopathy patients in this study was not available, we are confident that the clinical diagnoses are robust as current clinical practice is to test only individuals with a confirmed diagnosis *(15, 16)*. The proportion of cases with inherited cardiomyopathy is unknown, as evidence of familial disease is not a requirement for testing. The clinical sensitivity (proportion of patients with a pathogenic variant) in our case series was lower than earlier surveys that may reflect more restricted testing of stringently selected cases, typically from multiply-affected families with severe disease *(38, 39)*. However, the cohorts studied are representative of those encountered by clinical diagnostic laboratories, rather than a highly selected subset.

Despite high levels of confidence in interpreting many well-characterised variants (which may give ORs in the hundreds), diagnostic laboratories are understandably cautious when interpreting a variant that has not been seen before. Our analyses demonstrate that for many genes even variants currently reported as Uncertain Significance (VUS) show a several-fold case excess over the background in ExAC (Fig. 1). More refined interpretation of variants in validated genes, for example leveraging domain information, regional evolutionary constraint and cumulative clinical experience, could lead to substantial increases in the diagnostic yield of genetic testing, and indeed are likely to lead to much more substantial gains than the expansion of gene panels.

In contrast to the conservative strategy of clinical laboratories, research studies often report large “yields”. Some may not adequately control for the background rate of rare variation or may include genes for other conditions and as a result genes nominated as important contributors to disease in fact have little if any excess variation in cases. Testing of broad gene panels and overly inclusive interpretation of variants may lead to erroneous conclusions about pleiotropic effects of genetic variation *(29, 34)* and overestimates of double or compound mutations *(40, 41)* and the population prevalence of the disease if extrapolated from genetic variation *(42)*.

We highlight that despite the absence of demonstrable excess of overall rare variation in a gene, specific variants identified in family studies may still be disease-causing. However, if such variants are a small minority of rare variants in cases, clinical testing of that gene will yield more false than true positives. Moreover, for some of the genes that show no excess despite reasonable numbers of variants detected (Fig. 1) we note that the original reports did not include any variant with robust evidence of segregation (i.e. LOD >3) and here the possibility exists that the reported disease association is entirely spurious. An argument is often made that variants in candidate genes, even if not causal, could be contributing as modifiers *(43, 44)*. This remains possible but in the absence of any significant over-representation in cases the more parsimonious interpretation is that they are phenotypically silent. We have not tested more common variants (MAF>1×10^−4^), which could be mechanistically informative but are likely to have smaller effects *(45)* and have not evaluated individual level data to assess the impact of co-inheritance of variants, which are limitations of the analyses.

A further limitation of this study was the varying sequencing strategies in the case and control cohort, with the exome sequencing data of ExAC potentially less sensitive at detecting variants than clinical sequencing. We adjusted for this where coverage in ExAC is poor, while retaining only confidently-called variants. Additionally, if the frequency of rare variation in ExAC has still been marginally underestimated, it suggests that we have been conservative with respect to the key conclusions of this study, i.e. that variants in many previously associated genes are not enriched in cases. Another potential limitation of these analyses was that ethnicity data was not available for the OMGL or LMM HCM patients, therefore, we were unable to confirm the extent to which the cohorts used in this study were matched by race. However, by studying the aggregate burden of multiple very rare variants we expect that any confounding effects by individual population-specific variants in cases or controls will be limited. Supporting this assumption, an analysis of the LMM DCM cohort comparing findings from all populations with the Caucasian-only subsets revealed that the conclusions are robust.

Our findings highlight the need for systematic evaluation of the evidence of disease association and disease mechanisms (e.g. gain or loss of function) for clinical interpretation of putative disease genes, such as the ClinGen project (http://clinicalgenome.org *(46)*), alongside large population databases representing diverse ethnic origins, preferably linked to phenotypic data.

In conclusion, we have demonstrated that new opportunities for large-scale comparison of rare variation in Mendelian disease genes between patient cohorts and the wider population can highlight the genes, regions of genes and/or classes of variants which can be reliably interpreted in a clinical setting. For validated disease genes, there is clear potential to increase the yield of correctly interpreted, actionable variants. At the same time, problems must be avoided by recognizing that many implicated genes, and a significant proportion of variants, may not be robust. As clinical genetic testing moves to large gene panels and whole exome and genome sequencing, an understanding of gene and variant pathogenicity will be increasingly important in order to deliver reliable genomic interpretation.

## Materials and Methods

### Study Design

#### Clinical cohorts

##### Oxford Medical Genetics Laboratory (OMGL)

The OMGL cohort comprises apparently unrelated index cases referred from Clinical Genetics centers across the UK, with initial clinical diagnosis of HCM, DCM or ARVC made by a consultant cardiologist. All samples received for diagnostic genetic testing of HCM, DCM or ARVC genes were eligible and analysis was undertaken in a routine clinical setting using clinical consent. Comprehensive data on patient ethnicity is not available for this cohort. Genotype data was obtained from 3267 individuals with HCM, 559 with DCM and 361 with ARVC. The current panel sizes include 16 genes for HCM, 28 genes for DCM and 8 genes for ARVC. Not all patients were analyzed for every gene (see Supplementary Table S2). Variants reported in this cohort were classified according to national guidelines (http://www.acgs.uk.com) as highly likely to be pathogenic (Class 5), likely to be pathogenic (Class 4), or variant of unknown significance (VUS) (Class 3).

##### Laboratory of Molecular Medicine, Partners Healthcare (LMM)

Data from LMM was downloaded from the supplemental files of published HCM *(12)* (18 genes sequenced in 632 - 2912 patients) and DCM *(4)* (46 genes sequenced in 121 - 756 patients) cohorts. The LMM HCM cohort comprised unrelated probands referred for HCM clinical genetic testing. Any individuals with an unclear clinical diagnosis of HCM, or with left ventricular hypertrophy due to an identified syndrome such as Fabry or Danon disease, or unaffected individuals with a family history of HCM were excluded. The LMM DCM cohort comprised individual probands referred for DCM clinical genetic testing. According to the published report, all patients had DCM or clinical features consistent with DCM based on the medical and family history information provided by ordering providers. Additionally, any cases with confirmed diagnoses of other cardiomyopathies, structural heart disease, congenital heart disease or syndromic or environmental causes were not included in the study. Variants are classified as pathogenic, likely pathogenic, VUS favor pathogenic or others (other VUS, likely benign) according to the LMM’s clinical grade variant classification criteria *(4)*.

OMGL and LMM use similar clinical guidelines and employ equivalent approaches for variant classification in line with published guidelines *(5)*, however different names are used for each class. In this manuscript the term Pathogenic (P), includes OMGL Class 5, and LMM Pathogenic; Likely pathogenic (LP), includes OMGL Class 4 and LMM Likely pathogenic; and Variant of Uncertain significance (VUS), includes OMGL Class 3 and the LMM VUS favor pathogenic and other VUS (see Fig. 1).

Fisher’s exact test was performed using Stata statistical analysis software (V10.0) to compare the proportion of cases with rare (MAF 1×10^−4^) variants in each gene between the OMGL and LMM cohorts (see Supplementary Tables S3A, S3B). All statistical tests used in these analyses are two sided, unless otherwise stated.

In these clinical cohorts, sequence data was generated using a range of mutation scanning and direct sequencing techniques of varying sensitivity (High-resolution DNA melting, WAVE dHPLC, LightScanner^®^, DNA microarray [Cardiochip], Sanger sequencing, targeted Next Generation Sequencing [NGS]). These targeted tests are designed to cover the coding regions and splice sites of the key genes of interest; the analytical sensitivity of these methods is estimated to be in the region of 98-100% (data from in house validation). As all putative pathogenic variants are confirmed by Sanger sequencing the rate of false positive variant calls will be negligible.

### Exome Aggregation Consortium (ExAC) cohort

The ExAC dataset comprises aggregated sequencing data from a variety of large-scale exome sequencing projects, reprocessed through the same pipeline. VCF data was downloaded from the Exome Aggregation Consortium (ExAC), Cambridge, MA (http://exac.broadinstitute.org) [version 0.3, Jan 2015]. Quality control analyses suggest a sensitivity of 97-99.8% for single nucleotide variants (SNVs) and approximately 95% for insertions and deletions (indels)*(8)*. To minimize any bias resulting from the higher sensitivity strategies employed by diagnostic laboratories, only genes with a high proportion of coding region covered to a median sequence depth of >30× and only high quality (PASS filter) variants were included in our analyses. In addition we adjusted the total number of ExAC samples per gene based on the mean coverage at the variant sites of interest.

Sample size was fixed by the availability of clinically sequenced and ExAC samples. Illustrative power calculations are reported in Supplementary table S12. Table 1 provides an overview of all of the cohorts analyzed in this study.

**Table 1:** Overview of the datasets analyzed in this study. Clinical data is from the Oxford Medical Genetics Laboratories (OMGL), UK and the Laboratory of Molecular Medicine (LMM), USA (combined clinical datasets for HCM and DCM are shaded in grey).

| Dataset | Cohort Type | Reference | Sample size | No. genes tested | Methods used to generate sequence data |
| --- | --- | --- | --- | --- | --- |
| OMGL HCM | Clinical | this paper | 807 - 3267 | 16 | High-resolution DNA melting (WAVE dHPLC, LightScanner®)<br>Sanger sequencing<br>Targeted NGS |
| LMM HCM | Clinical | (12) | 632 - 2912 | 18 | DNA microarray (Cardiochip)<br>Sanger sequencing.<br>Targeted NGS |
| Combined HCM | Clinical | - | 632 - 6179 | 20 | As detailed above |
| OMGL DCM | Clinical | this paper | 304 - 559 | 28 | High-resolution DNA melting (WAVE dHPLC, LightScanner®)<br>Sanger sequencing<br>Targeted NGS |
| LMM DCM | Clinical | (4) | 121 - 756 | 46 | DNA microarray (Cardiochip)<br>Sanger sequencing<br>Targeted NGS |
| Combined DCM | Clinical | - | 121 - 1315 | 48 | As detailed above |
| OMGL ARVC | Clinical | this paper | 93 - 361 | 8 | High-resolution DNA melting (WAVE dHPLC, LightScanner®)<br>Sanger sequencing<br>Targeted NGS |
| Research HCM | Research | (34) | 874 | 36 | Targeted NGS |
| Research DCM | Research | (30, 35, 36) | 312 - 324 | 12 | Sanger sequencing |
| ExAC | Database | http://exac.broadinstitute.org | Up to 60706 | all | Exome sequencing |

### Calculation of frequency of rare variation in cardiomyopathy cohorts and ExAC

A minor allele frequency (MAF) cut off of 1×10^−4^ was used to define rare variants – the Results section and Supplementary Note 1 describe how this was defined. For each gene, the overall frequency of ExAC variants with a minor allelic frequency (MAF) below the selected threshold was calculated by dividing the sum of the adjusted allele count by the mean of the total adjusted alleles. Only likely protein-altering variants in designated canonical transcripts (Supplementary Table S2) were analyzed, i.e. missense, in-frame insertions/deletions, frameshift, nonsense and variants affecting the splice donor and acceptor regions (first and last two bases of each intron). Analyses were performed on all protein-altering variants and separately on variants predicted to be non-truncating (missense and in frame insertions and deletions) and variants predicted to be truncating (frameshift, nonsense, splice donor/acceptor).

The frequency of rare variation in the cardiomyopathy cohorts was calculated by dividing the sum of rare variants identified in cardiomyopathy cases by the total number of patients analyzed for each gene. In total, after combining data from both clinical laboratories and excluding poorly covered genes in ExAC, 20 genes sequenced in 632 - 6179 HCM patients, 46 genes sequenced in 121 - 1315 DCM patients and 8 genes sequenced in 93 - 361 ARVC patients were analyzed. See Supplementary Table S2 for full details of cohort sizes for each gene.

### Comparison of variation between disease cohorts and ExAC controls

For each gene, the frequency of rare variation observed in the clinical cohort was compared to that observed in ExAC. The case excess was defined by subtracting the proportion of individuals in ExAC with a filtered variant, from the proportion in the clinical cohort. For these analyses we made the simplifying assumption that the frequency of rare, benign variants was equivalent in cases and controls and that the frequency of pathogenic variants in the ExAC is sufficiently low so as not to affect this comparison (as the highest estimated prevalence of these diseases are 1 in 500 people*(13)*. Fisher’s exact test was performed using Stata software. The level of significance, p=0.05, was adjusted with Bonferroni correction for each gene set (HCM P<0.0025, DCM P<0.001, ARVC P<0.006).

For each gene and variant class we calculated two related metrics: the odds ratio (OR) and the etiological fraction (EF)*(25–27)* (for further information on EF please refer to Supplementary Note 2). ORs (with 95% confidence intervals) were calculated using Stata software. For cells with zero values a 0.5 correction was added to all cells before calculating the OR. The EF was calculated as follows: (OR-1)/OR × 100. EF values and ORs were calculated for all protein-altering variants and separately on predicted non-truncating and predicted truncating subsets.

Figures summarizing statistical analyses are based on binary (case/control) methods that estimate odds ratios, these methods are assumption-free with respect to data distributions.

### Assessment of population-specific effects

As ethnicity data was not available for the OMGL cases or for the specific variants identified in the LMM HCM cohort, the total case and control datasets were used in this study. ExAC is a mixed dataset but with a majority of samples of European descent (52% non-Finnish European) which should be well matched to the LMM cohorts (62% white/Caucasian) and the unselected UK referral population of the OMGL cohorts. To assess if population-specific variants in these full datasets had any confounding effects, we compared the results observed between the LMM DCM cohort (for which individual self-reported patient ethnicity data is available) and ExAC for the full datasets and for white DCM patients (456 samples) versus the ExAC non-Finnish European subset (33,370 samples). Discrepancies were defined as any gene which had a significant case excess (as defined above) in one of the analyses but not the other. The correlation (Pearson coefficient) between case excess frequencies observed in both analyses was also calculated for genes with more than 300 samples sequenced.

### Distribution of missense variants in MYH7

To identify putative hotpots of pathogenic missense mutations in *MYH7*, distinct rare missense variants in the clinical HCM and DCM cohorts and the ExAC population controls were mapped along the protein sequence. Non-random mutation cluster (NMC)*(47)*, implemented in the iPAC Bioconductor R package, was used to identify clusters of variants in each cohort. (R source code of NMC algorithm: https://www.bioconductor.org/packages/devel/bioc/html/iPAC.html)

### Analysis of research cardiomyopathy cohorts

Research cardiomyopathy cohorts are defined as published studies from research laboratories where patient samples were subjected to sequencing across panels of cardiac genes, but for which clinical grade variant classification was not performed. Instead, putative pathogenic variants were defined based on the type of variant (truncating or other), presence or absence in small control or population cohorts and databases and the output of *in silico* algorithms used to predict the effect of missense variants. The full details of the classification criteria used in each study are described in the references described below.

The HCM research cohort*(34)* sequenced 874 patients across 35 genes (12 primarily associated with HCM, 7 with DCM, 7 with ARVC and 9 with arrhythmias, as stated by Lopes et al.). The DCM research cohort*(30, 35, 36)* comprised 312 - 324 patients sequenced for 12 confirmed and putative DCM genes (not including TTN). For all cohorts, variant lists were downloaded and rare variant frequencies and case excess were calculated for each gene as described for the clinical cohorts above.

To assess the accuracy of the variant classification methods used in these published research cohorts, the number of variants reported to be putatively pathogenic for each gene in these studies was compared to the number predicted to be pathogenic based on the case excess observed in these cohorts.

### Coverage of HGMD cardiomyopathy mutations in ExAC

Variants in the Human Genome Mutation Database (HGMD, professional version 2015.1) associated with HCM, DCM or ARVC were identified based on manual curation of the HGMD disease terms. Only HGMD “disease-causing mutations” were assessed - analysis was performed on all variants (with a HGMD tag of DM and DM?) and separately on just those variants with a DM tag (according to HGMD documentation, variants with a tag of DM? have a degree of doubt with regard to pathogenicity). The total allele frequency and adjusted allele count from ExAC was extracted for each variant. Polymorphisms (defined as an ExAC allele frequency >1×10^−2^) were removed from the analysis. The number of HGMD variants present in ExAC was calculated, at any frequency and at a common frequency (MAF > 1 × 10^−4^) highly unlikely to be compatible with pathogenicity. The total number of ExAC alleles and the total number of ExAC individuals with HGMD-associated cardiomyopathy variants was also calculated for each disease. Additionally, the ExAC frequencies of HGMD cardiomyopathy variants previously observed only once in the Exome Sequencing Project (ESP; http://evs.gs.washington.edu/EVS/) were analyzed to assess how the enhanced resolution of ExAC can clarify previously uninterruptable variants.

## Supplementary Materials

Supplementary Note 1: Selecting an allele frequency threshold to define potentially pathogenic and penetrant variants in Mendelian conditions

Supplementary Note 2: Etiological fraction (EF)

Fig. S1. Comparison of the excess of rare variants in LMM DCM cases over controls between all population analysis and Caucasians only.

Table S1A. Variants identified in HCM, DCM and ARVC patients tested at Oxford Medical Genetics Laboratories (OMGL).

Table S1B. Variants identified in HCM and DCM patients tested at Partners Healthcare Laboratory of Molecular Medicine (LMM)

Table S2. Genes and transcripts analysed in this study and the number of patients sequenced in each disease cohort.

Table S3A. Comparison of the frequency of pathogenic variants in tested genes between OMGL and LMM clinical laboratories for HCM cohorts

Table S3B. Comparison of the frequency of pathogenic variants in tested genes between OMGL and LMM clinical laboratories for DCM cohorts

Table S4A. Comparison of the frequency of rare variation (ExAC MAF < 0.0001) in clinical HCM cases compared to ExAC controls.

Table S4B. Comparison of the frequency of rare variation (ExAC MAF < 0.0001) in clinical DCM cases compared to ExAC controls.

Table S4C. Comparison of the frequency of rare variation (ExAC MAF < 0.0001) in clinical ARVC cases compared to ExAC controls.

Table S5A. Odds ratios and Fisher’s Exact test results testing for significance of the excess of rare variation in HCM cases versus ExAC controls.

Table S5B. Odds ratios and Fisher’s Exact test results testing for significance of the excess of rare variation in DCM cases versus ExAC controls.

Table S5C. Odds ratios and Fisher’s Exact test results testing for significance of the excess of rare variation in ARVC cases versus ExAC controls.

Table S6. Frequency of rare variants in HCM research cohort with comparison between the number of variants reported as pathogenic by Lopes et al with the number predicted by the case excess observed versus ExAC controls.

Table S7. Frequency of rare variants in DCM research cohort with comparison between the number of variants reported as pathogenic by Hershberger et al with the number predicted by the case excess observed versus ExAC controls.

Table S8. Numbers of variants in the HGMD database (professional version 2015.1) associated with HCM by gene with number of these variants present in ExAC (at any frequency and greater than 0.0001), the Exome Sequencing Project (ESP) or 1000 Genomes (1KG) and the total number of ExAC alleles and individuals with a HCM-associated variant.

Table S9. Numbers of variants in the HGMD database (professional version 2015.1) associated with DCM by gene with number of these variants present in ExAC (at any frequency and greater than 0.0001), the Exome Sequencing Project (ESP) or 1000 Genomes (1KG) and the total number of ExAC alleles and individuals with a DCM-associated variant.

Table S10. Numbers of variants in the HGMD database (professional version 2015.1) associated with ARVC by gene with number of these variants present in ExAC (at any frequency and greater than 0.0001), the Exome Sequencing Project (ESP) or 1000 Genomes (1KG) and the total number of ExAC alleles and individuals with an ARVC-associated variant.

Table S11. Summary of the numbers of variants in the HGMD database (professional version 2015.1) associated with HCM, DCM and ARVC which are present in ExAC (at any frequency and greater than 0.0001), the Exome Sequencing Project (ESP) or 1000 Genomes (1KG) and the total number of ExAC alleles and individuals with a ARVC-associated variant.

Table S12. Power calculations.

Table S13. Comparison of case-control analysis for LMM DCM cases with (1) Caucasian DCM patients versus ExAC Non-Finnish European subset and (2) All DCM patients versus total ExAC dataset.

## Acknowledgments

We thank the staff at the Oxford Medical Genetics Laboratories (OMGL), Oxford University Hospitals NHS Foundation Trust, for generating and interpreting data used in these analyses.

## Author contributions

HCW, SAC, MF, RW, KLT and JSW designed the study, analysed and interpreted the data, and wrote the manuscript. FM, EVM, DGM, JW, KLM, EB, AS, JCT and ExAC generated and analysed data. All authors approved the final manuscript.

**Competing interests**: None

**Data and materials availability**: OMGL ClinVar Database Accession No: [Placeholder].

**Figure S1:**
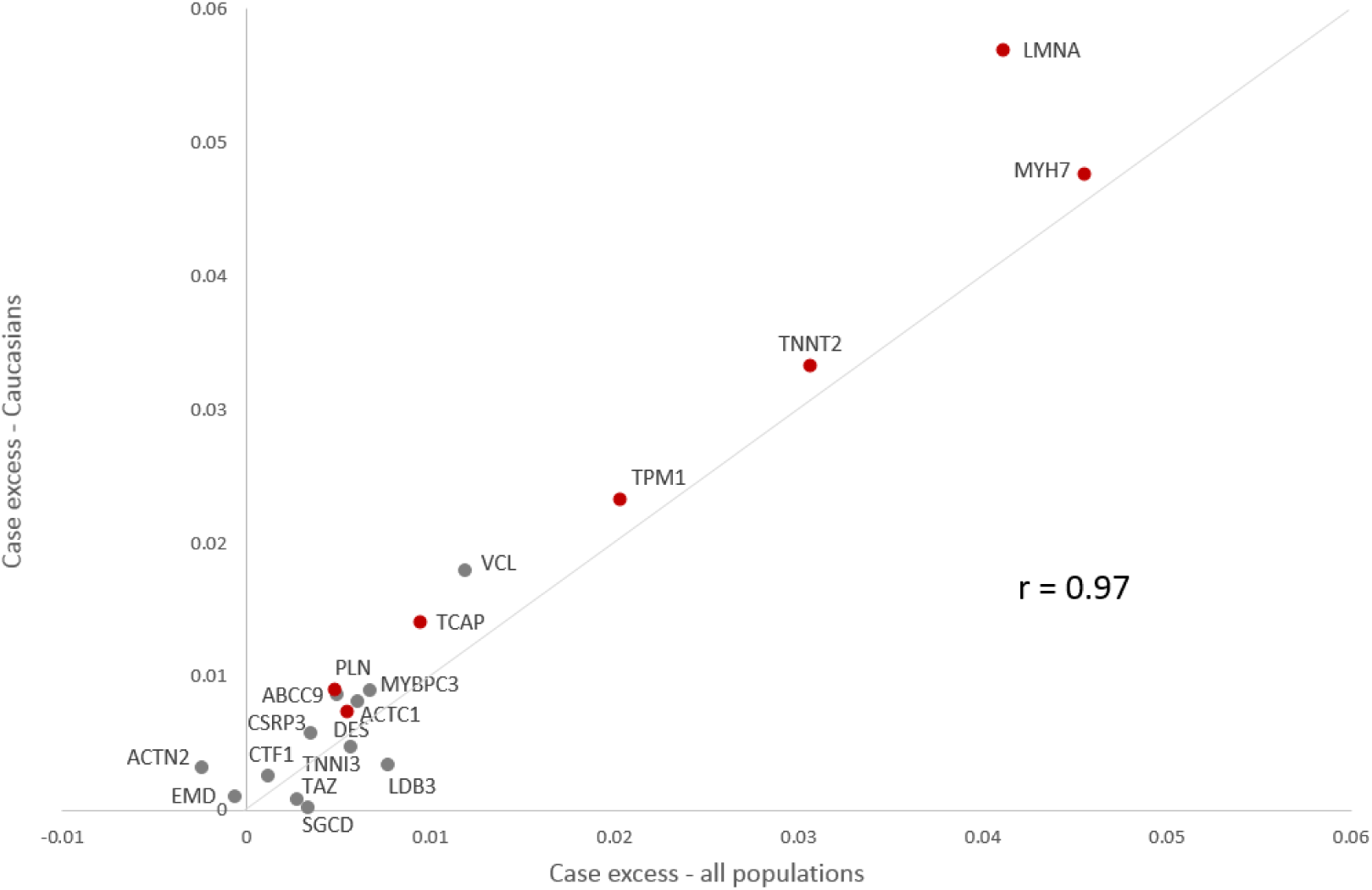
Comparison of the excess of rare variants in LMM DCM cases over controls between all population analysis (all cases versus full ExAC dataset) and Caucasian only (Caucasian DCM cases versus non-Finnish European subset of ExAC). Genes with a significant excess in cases (Fisher’s exact test p<0.001 with Bonferroni correction for 48 tests) are shown in red. The Pearson correlation coefficient was calculated at 0.97. Only genes with >300 samples sequenced are shown.

**Supplementary Note 1**

*Selecting an allele frequency threshold to define potentially pathogenic and penetrant variants in Mendelian conditions*

How many times must a variant be observed in ExAC to consider it too common to cause a Mendelian cardiomyopathy?

Clearly, a penetrant Mendelian allele should not be present in an unselected population more frequently than the disease it causes. Moreover, for a genetically heterogeneous condition it must not be more frequent than the proportion of cases attributable to that gene, or indeed to any single variant. Given the substantial datasets presented here we can now estimate these proportions reasonably robustly, but must take care to provide an appropriate margin of error given that these estimates are derived from samples of fixed size and ethnicity.

HCM has an estimated prevalence of 1:500*(13)*. In this series the variant to which the largest proportion of cases is attributable is MYBPC3 c.1504C>T (p.Arg502Trp), found in 104/6179 HCM cases (1.7%, 95CI 1.4-2.0%). Caution must be applied when considering calculating confidence intervals for allele frequencies, especially for very rare alleles, as the underlying distribution of allele frequencies from which our sample is drawn is not fully known, and strongly left skewed (e.g. a singleton variant is much more likely to have a true frequency below the measured frequency than above it). Nonetheless, the binomial distribution is a fair approximation at this allele frequency range, and will generally be conservative when calculating upper confidence intervals for the frequency of rare alleles, though not at all robust for calculating lower confidence intervals.

Given that MYBPC3c.1504C>T is seen in 1.7% of cases (in the heterozygous state), and cases have a prevalence of 1:500, we expect a population allele frequency around 1.7 ×10-5. 104/6179 × 1/500 × 1/2 (as each individual is diploid) = 1.7×10-5

We can cross-reference this against ExAC. This variant is observed 3 times in ExAC (in 60557 individuals genotyped at this site), and twice in 33329 Europeans (non-Finnish). This gives an observed ExAC global MAF = 2.5×10-5 (6.0×10-5 in Europeans), and an upper bound for the true population frequency of this allele would be estimated as 5.3 x10-5 based on a binomial distribution around the ExAC global MAF, or 7.2×10-5 based on the European subset.

*# upper limit based on observation in global ExAC*

binom::**binom.confint**(3,2*60557,methods="asymptotic",confint=0.95)$upper

\## [1] 5.279914e-05

*#upper limit based on observation in ExAC Europeans*

binom::**binom.confint**(2,2*33329,methods="asymptotic",confint=0.95)$upper

\## [1] 7.15858e-05

So we consider that a variant with a true population allele frequency > 7 × 10-5 as too common to cause HCM. We may want to adjust this threshold to allow for reduced penetrance. We could simply divide this maximum frequency by the penetrance of the condition (e.g. a variant with a penetrance of 0.5 could be present at double this frequency).

In fact we have made a series of conservative assumptions (and more are to come), so have not made an additional correction for reduced penetrance at this point.

Finally, we ask how many times such a variant might be a variant with true population allele frequency 7×10-5 be observed in a random population sample. This can be modeled using a poisson distribution: for a 5% error rate we take the 95th centile of a poisson distribution with λ = expected allele count given by 2 × sample size × population allele frequency:

maxAF=7E-5

myCI=0.95

nSamples=60706

alleleNumber=2*nSamples

maxAC = **qpois**(myCI, alleleNumber*maxAF)

maxAC ## [1] 14

This allows us to adjust our threshold for variants that are not sequenced in the entire cohort (indicated by the “AN” field in ExAC), or by focusing on one ethnic subgroup. For example, if the variant was found only in East Asians, we could derive an ExAC AC cut-off as follows:

nSamples=4327 *# (ExAC EAS)*

**qpois**(myCI,2*nSamples*maxAF)

\## [1] 2

**Limitations**

In the populations studied, a measured AF greater than this threshold is incompatible with penetrant Mendelian cardiomyopathy. However, deleterious founder variants may be present in other populations in whom the genetic architecture of CM is undefined. As described here, we have not accounted for reduced penetrance.

The maximum allele frequency that is compatible with pathogenicity will be different for each condition according to its prevalence and the known genetic architecture of the disease.

Examples here are generated using the R statistical environment.

**Supplementary Note 2**

*Etiological fraction (EF)*

*Attributable risk percent among exposed* (ARP), provides an estimate of the proportion of the risk in an exposed population that can be attributed to the exposure. This epidemiological measure is meaningful where there is strong evidence of a biologically plausible causal relationship between exposure and disease. In our context, ARP corresponds to the proportion of the risk of cardiomyopathy in mutation carriers that can be attributable to the mutation. The measure was popularized by Cole % MacMahon *(25)*, and defined as follows:

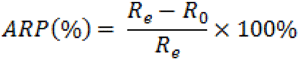

where

***R*_*e*_** = risk in the exposed (i.e. carriers of rare variant in gene of interest)

***R*_*a*_** = risk in the unexposed (i.e. individuals with no rare variant in gene of interest)

The equation can be conveniently rewritten as:

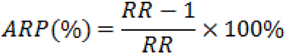

where

***RR*** = relative risk (ratio of risk among exposed to risk among unexposed) (Robins % Greenland *(27))*

For cross-sectional (i.e. case-control) data, odds ratios (OR) provide accurate estimates of the underlying relative risk *(49)*, leading to:

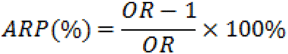

Now ARP, when expressed as a decimal fraction, has been called the etiological fraction (EF) *(27)* i.e.

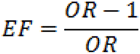

and we refer to EF hence-with.

In the diagnostic context, where we are treating cardiomyopathy as a Mendelian disease, and are interested in interpreting whether an individual rare variant was likely causative, the EF can be interpreted in several ways^**a**^:

- the proportion of variant-carrying probands in which the variant was causal
- the proportion of variants, found in affected individuals, that were penetrant
- the probability that an individual rare variant, found in a proband, was responsible for the disease

EF therefore represents our confidence in interpreting variation as etiologically significant when found in an individual with disease.

## Footnotes

a It is worth noting that in each case we cannot disentangle incomplete penetrance. Without additional information we cannot determine whether an EF of 50% indicates that half of the variants found in our case cohort are fully penetrant and disease-causing (and the other half have zero penetrance), or if all of the variants are pathogenic, but with reduced (50%) penetrance.

